# Debiasing Sinkhorn divergence in optimal transport of cellular dynamics

**DOI:** 10.1101/2025.01.11.632566

**Authors:** Jayshawn Cooper, Christina Young, Pilhwa Lee

**Affiliations:** Department of Mathematics, Morgan State University, 1700 E. Cold Spring Lane, Baltimore, 21251, MD, USA

**Keywords:** algorithmic bias, optimal transport, single cell analysis, entropic regularization, Sinkhorn divergence, lineage determination

## Abstract

Single-cell RNA-seq analysis characterizes developmental mechanisms of cellular differentiation, lineage determination, and reprogramming with differential conditioning of the microenvironment. In this article, the underlying dynamics are formulated via optimal transport with algorithms that calculate the transition probability of the state of cell dynamics over time. The algorithmic biases of optimal transport (OT) due to entropic regularization are balanced by Sinkhorn divergence, which normally de-biases the regularized transport by centering them. In the case of reprogramming mouse embryonic fibroblasts [1] with dense time points, Sinkhorn divergence is shown to improve the trajectories of targeted cell fates depending on the specific cell types. When the time points are filtered out with sparser 9 and 5 time points, some cell phenotypes show better outcomes from strong entropic regularization. For 9 time points with 2 days interval, Sinkhorn divergence shows a clear advantage with broad bandwith of optimal entropic regularization. For these derived time points, when the cell population is scaled down from ***n* = 8000** to ***n* = 2000**, there comes no benefit from Sinkhorn divergence for some specific cell types. In the case of stratifying morphogenesis of the epidermis [2], the sparsity of time points makes it not significant to prescribe Sinkhorn divergence in the accuracy of transporting to the expected cell fates. Overall, whether to prescribe Sinkhorn divergence for the accurate prediction of lineages of single cells depends on temporal sparsity.

## 1 Introduction

Single-cell analysis illuminates the heterogeneity of genetic and epigenetic signatures in cellular distributions crucial in development and disease. Characterizing their temporal dynamics is significant in deconvolving the hidden nature of physiological mechanisms and therapeutic potentials [3]. Representing trajectories of cellular differentiation and programming/reprogramming is non-trivial due to the heterogeneity, and one rigorous approach is applying probability distribution of their gene expressions. Accordingly, it is available to represent a phenotypic transition from one state to another via optimal transport, i.e., transiting probability measure in the principle of minimal least action. There have been several algorithms for optimal transport for single-cell analysis. In the Eulerian point of view, Waddington-OT is based on the dual formulation of Kantorovich in Wasserstein distance with empirical source and target distributions prescribed as constrained optimization (unbalanced optimal transport) augmented with entropic regularization [1, 4]. Also, in the aspect of Lagrangian point of view of cellular trajectories, probability density can be estimated by Neural-ODE and normalized flow [5]. Recently, state-of-the-art techniques have been developed such as applying autoencoder with latent spaces [6], Gromov-Wasserstein for further resolving *in situ* loci [7], and predicting perturbation responses [8] among many others.

Taking optimal transport with empirical finite measures is mostly on non-convex and non-differentiable distributions. That is how the diffusive term is prescribed as entropic regularization transforming the optimzation problem globally convex [9]. In high-dimensional distributions with tens of thousands of RNA transcripts in a single cell, scalable algorithms make the analysis practical [10, 11]. Even though it is necessary to prescribe entropic regularization for improving optimal transport, the regularization does induce an algorithmic bias. One primary remedy of the bias induced by entropic regularization is Sinkhorn divergence [12]. There is one study investigating the functionality of measuring cell similarities from single-cell analysis via optimal transport with Sinkhorn divergence, i.e. debiasing Sinkhorn divergence [13]. Interestingly, prescribing Sinkhorn divergence does not guarantee reducing algorithmic bias due to entropic regularization. There are strong finite-sample effects of no improvement when entropic regularization is weak [14].

In this article, we explore whether debiasing Sinkhorn divergence improves the accuracy of generating expected trajectories of cell lineages and minimizing dispersed unexpected cell phenotypes. We assumed the main criteria for this is the sparsity of time points in single-cell datasets, i.e., the presence of strong nonlinearity in transporting from one distribution to another. Accordingly, two case studies were performed; one is about fibroblast reprogramming which is densely collected twice a day for 18 days [1]. The other is epidermal stratifying morphogenesis which is sparse only with two time points of Embryonic Days 12.5 and 17.5, i.e. E12.5 and E17.5 [2]. When time points are dense, Sinkhorn divergence with weak entropic regularization does improve the accuracy of cell fates for some specific cell types, but possibly not all. To investigate the effects of sparsity in time, the dataset of fibroblast reprogramming was filtered out in time points to 9 and 5, i.e., the time intervals to 2 and 4 days. Accordingly, the influence of the scale of population is also investigated by downsizing the cell population from *n* = 8000 to *n* = 2000. In the cases where time points are limited with their sparsity, sometimes there needs a strong entropic regularization depending on specific cell phenotypes.

## 2 Mathematical formulation

We represent cellular states in the probability distribution. Accordingly, the similarities between two cellular states are quantified by the distance between their probability distributions on Wasserstein metric spaces.

### 2.1 Wasserstein distance

When Wasserstein distance is considered for the distance between two probability measures *µ* and *ν* on Ω ⊆ ℝ^*n*^, the original formalism is based on the Monge map [15],

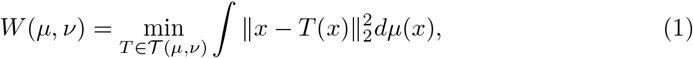

where

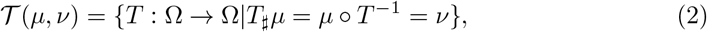

and *T*_*♯*_*µ* is called pushforward of *µ* through the Monge map *T*. Kantorovich has generalized the admissibility of Wasserstein distance in the following [16]:

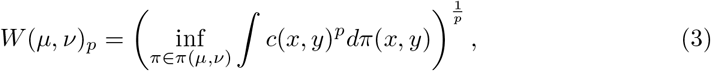

where *c*(*x, y*) is the cost function programming a cell at the state *x* to the state *y*, and the states are represented by RNA gene expression in our cases. As an element of the space of joint distributions of *π*(*µ, ν*), *π* is the joint distribution of the paired state (*x, y*) with its projection to *µ* and *ν* as marginal distributions.

### 2.2 Dynamic optimal transport

Dynamic optimal transport (OT) represents the Wasserstein distance from the view-point of dynamical systems. The *L*^2^-Wasserstein distance can be measured by the dynamic OT with the following formula [17]:

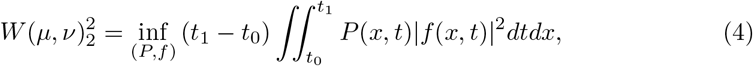

where *µ* and *ν* are two distributions at timepoints *t*_0_ and *t*_1_ and *f* (*x, t*) represents the velocity or drift term at the state *x* and time *t*, i.e., 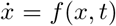. The probability distribution *P* (*x, t*) is evolved by the Liouville equation [18]:

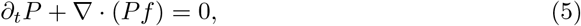

with the initial and final conditions prescribed by *P* (·, *t*_0_) = *µ* and *P* (·, *t*_1_) = *ν*.

### 2.3 Unbalanced optimal transport with entropic regularization

Among the aforementioned formulations for optimal transport and Wasserstein distances, the Waddington OT is an algorithm based on the dual formulation of Kantorovich for computing trajectory probabilities at specific times [1]. For a given set of cells *C* at time *t*_*j*_. The probability *p*_*tj*_ is defined as follows:

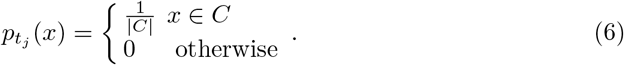

The descendant distribution at time *t*_*j*+1_ is calculated by *pushing* the cell set through the transport/cost matrix. Each probability is *pushed forward* by multiplying by the transport map on the right:

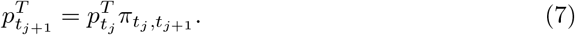

Therefore, inductively the descendant distribution can be calculated at any later time *t. > t*_*j*_. Waddington-OT’s approach employs both entropic regularization and unbalanced transport to compute the transport map at time *t*_*i*_ and *t*_*i*+1_. They solve the following optimization problem [1]:

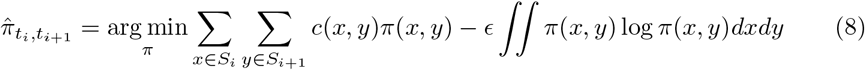

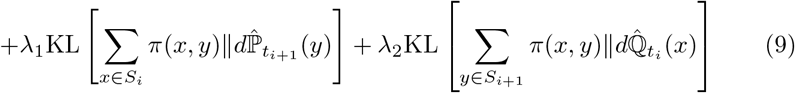

where *ϵ, λ*_1_ and *λ*_2_ are regularization parameters. 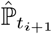 and 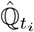 define the probability distributions from the empirical developmental process. 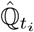 is from rescaling 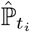 by the relative growth rate. *S*_*i*_ is a set of expression profiles at time *t*_*i*_. Computing the ancestors of *C* at an earlier time point *t*_*i*_ *< t*_*j*_ is a similar process, except the cell set is “pulled back” through the transport matrix:

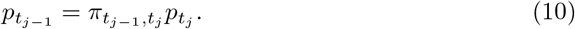

Having an ancestor distribution at an earlier time point and a descendant distribution at a later time point defines the *trajectory* for a cell set *C* [1].

### 2.4 Debiasing with Sinkhorn divergence

The entropic optimal transport is with the following objective function [14]:

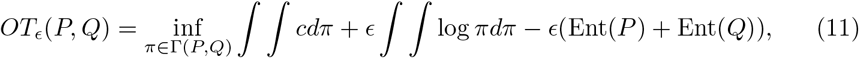

where the entropy of an absolutely continuous probability measure is denoted by Ent(*P*) = ∫*p*(*x*) log(*p*(*x*))*dx*, and *OT*_*ϵ*_(*P, Q*) is strongly convex to provide a unique optimality. Waddington-OT retaining the entropic optimal transport tends to lead to algorithmic bias by altering a cell’s transport distance. Therefore, when computing transport matrices for the Waddington-OT pipeline, we utilize *Sinkhorn Divergence* to compute alternative and debiased transport matrices for each pair of timepoints, i.e., Sinkhorn divergence is a centering method that can be used to debias the algorithm [14]:

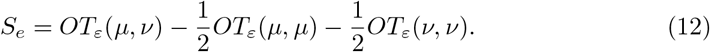

such that *S*_*ε*_(*P, Q*) = 0 ↔ *P* = *Q*.

### 2.5 Comparison of the empirical ground-truth distribution and trajectories from optimal transport

In order to obtain insight into the overall impact of the entropy parameter in Eq. (8), we developed an algorithmic pipeline to determine the Wasserstein distance between the probability trajectories of each phenotype against the ground-truth (empirical) distribution of the cells prescribed by both Entropic regularization (Entropic map) and Sinkhorn divergence (Sinkhorn map) with in the interval from 0.01 to 2. On the final day of *t*_FINAL_, the ground truth distribution, *p*_ground_truth_ is represented following Eq. (6) where *C* is the subpopulation of the targeted cell phenotype. The Wasserstein distance *W* (*p*_ground_truth_, *p*_*t*FINAL_) indicates the accuracy of trajectory probability quantified by the distance to the targeted cell type, where *p*_*t*FINAL_ is from Eq. (7).

### 2.6 The theoretical basis for the counter-examples against outperformance of Sinkhorn divergence

The existence of entropic parameters degrading the performance of Sinkhorn divergence is addressed in [14]. The following Brenier’s Theorem, Kantorovich and self potentials are a brief exposition for Theorem 2 summarizing those in [14].

For Ω ⊆ ℝ^*d*^ compact, we define 𝒫(Ω) to be the space of (Borel) probability measures with support contained in Ω, and 𝒫_ac_ be those with densities.

#### Theorem 1

(Brenier’s Theorem(1991) [19]). *Let P* ∈ 𝒫_ac_(Ω) *and Q* ∈ 𝒫(Ω). *Then there exists a solution T*_0_ *to Eq. (1), with T*_0_ = *Id* − ∇*f*_0_ *where f*_0_ *is a 1-strongly concave function solving*

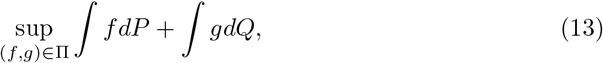

*where* 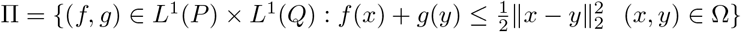.

The maximizers (*f*_0_, *g*_0_) to Eq. (13) are *optimal (Kantorovich) potentials*. The Entropic OT problem admits the following dual formulation under mild conditions,

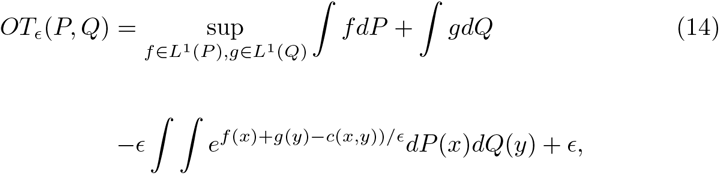

where the maximizer, denoted (*f*_*ϵ*_, *g*_*ϵ*_) are called *optimal entropic potentials*. We denote by *α*_*ϵ*_ the optimal entropic *self-potential* :

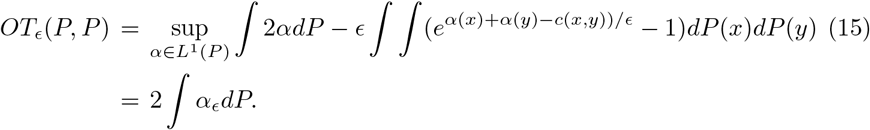

For a given *T*_0_ and any estimator *S* : ℝ^*d*^ → ℝ^*d*^,

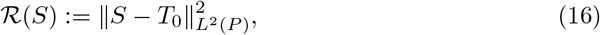

where 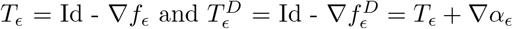.

#### Theorem 2

(Pooladian, Cuturi, and Niles-Weed (2022) [14]). *For any ϵ* < 1 *and any M >* 0, *there exist a pair of densities* (*P*_*ϵ*_, *Q*_*ϵ*_) *for which* 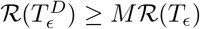.

## 3 Numerical implementation

The unbalanced optimal transport is taken from Waddington-OT [1]. We utilize Sinkhorn Divergence provided by the Python-OT package [20] to compute alternative and debiased transport matrices for each pair of timepoints. For computing the Wasserstein distances for the accuracy of trajectories prescribed by entropic regularization or Sinkhorn divergence in comparison to the probabilistic distribution of the ground truth phenotypes, the function *emd2* provided by the Python-OT package is employed. For the convergence of Sinkhorn divergence, the maximum iteration was set to 20000.

## 4 Results and Discussion

The main results are from two case studies, fibroblast reprogramming, which is dense in the time points with 18 days [1], versus stratification of epidermal cells in morphogenesis, which is sparse in time points only with 2 days [2].

### 4.1 Case Study 1: Fibroblast reprogramming

From a dataset of 315,000 cells collected over an 18-day period at half-day intervals, the modified Waddington-OT algorithm with Sinkhorn divergence is applied to a subset of 8,000 cells. The cells at Day 1 are transited by optimal transport in the expectation of reprogramming them to stromal, iPS, trophoblast, and epithelial cells as shown in Fig. 1(B). One track is through the trajectories with entropic regularization. The other track is through the trajectories with entropic regularization balanced by Sinkhorn divergence. For each cell type, the influence of entropic regularization is determined by *ϵ*. Accordingly, we have quantified the accuracy of reaching at targetted phenotypes by Wasserstein distance between the probability distributions of the ground truth from experimental data and the derived phenotypes from the trajectories of optimal transport in terms of *ϵ* (Fig. 3). The probability distributions through the trajectories of stromal, induced pluripotent, trophoblastic, and epithelial cells at Day 12 are shown in Figures 4, 5, 6, and 7, respectively. The four subpanels in each figure show the combination of *ϵ* of 0.05 or 1.0 and the entropic regularization with and without Sinkhorn divergence. Consistent with Theorem 2, for trophoblast and epithelial cell types, small entropic parameters do not guarantee the outperformance of Sinkhorn divergence.

**Fig. 1.**
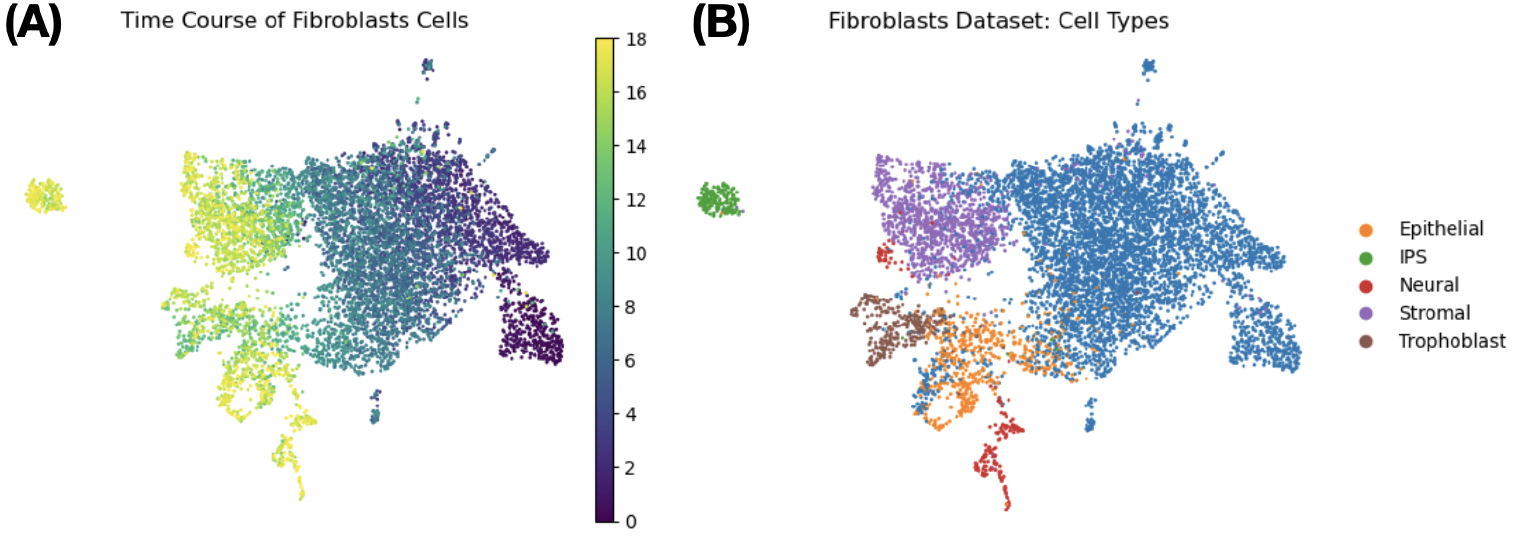
Reprogramming of fibroblasts over the 18-day period. (A) The heat map of reprogramming times from Day 1 to Day 18 with half-day intervals. (B) The reprogrammed lineages are represented by stromal, neural, iPS, trophoblast, and epithelial phenotypes [1].

**Fig. 2.**
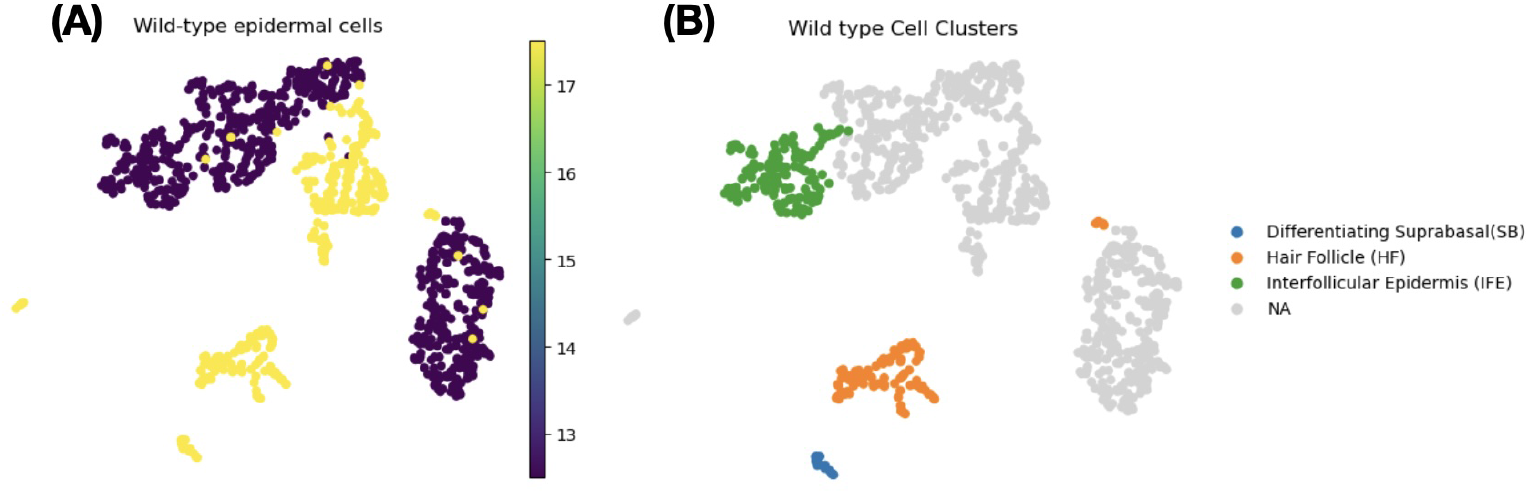
Stratification of mouse epidermal tissue in morphogenesis. (A) The heat map of two time points, E12.5 and E17.5. (B) The epidermal lineages are represented by suprabasal, hair follicle, and interfollicular epidermis phenotypes [2].

**Fig. 3.**
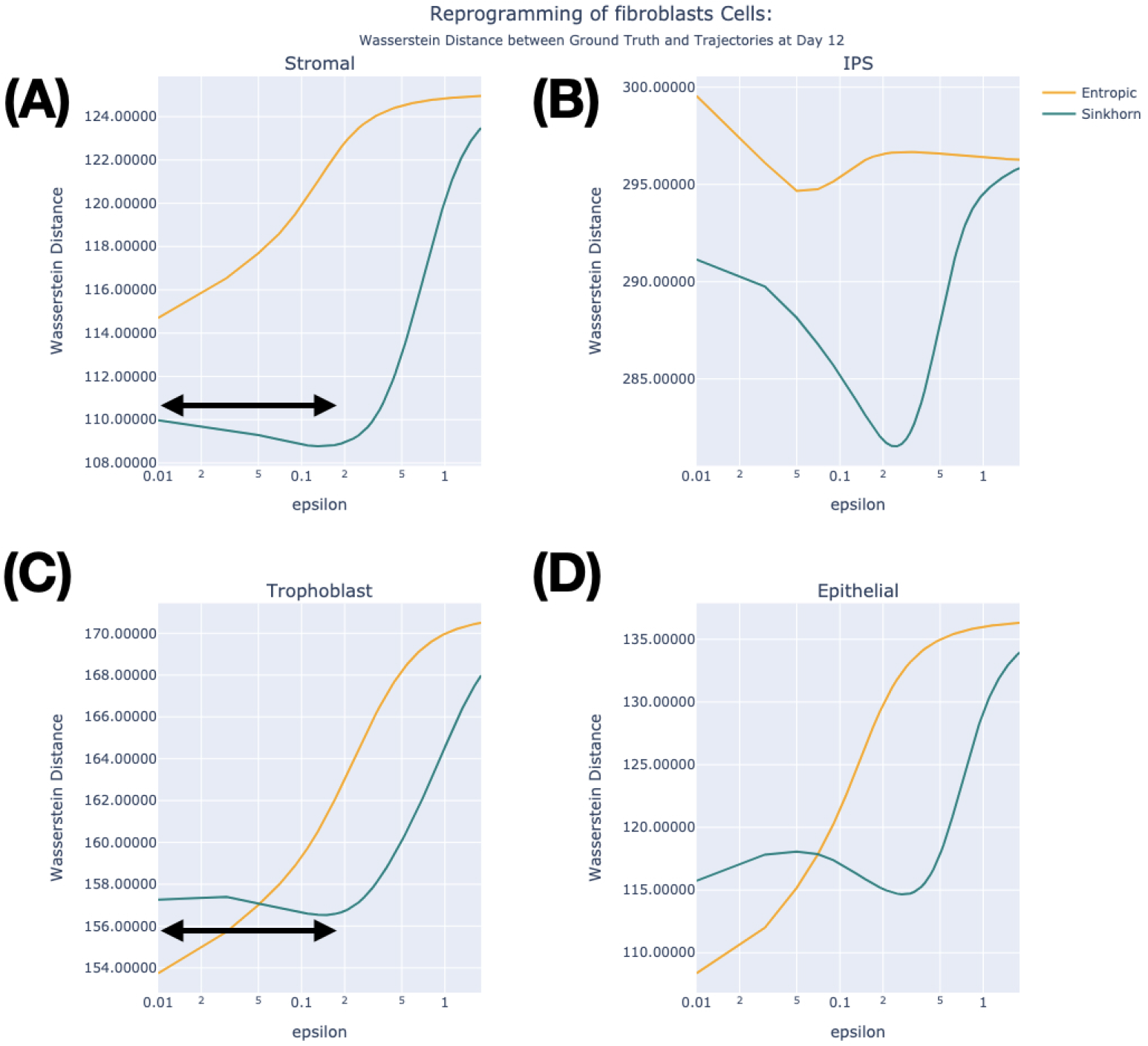
Accuracy of optimal transport for fibroblast reprogramming. Wasserstein distances between the probabilities of the ground truth phenotype and the trajectories of the entropic regularization and debiasing Sinkhorn divergence are plotted in terms of *ϵ*. Each panel is for stromal, iPS, trophoblast, and epithelial phenotypes. For stromal and iPS targeted trajectories, Sinkhorn divergence shows consistent improvement independent of entropic parameter *ϵ*. For trophoblast and epithelium targeted trajectories, the entropic map is better when *ϵ* is small. For stromal and trophoblast cells, competent performance of Sinkhorn divergence has a tolerated bandwidth of *ϵ* as indicated by the black arrows.

**Fig. 4.**
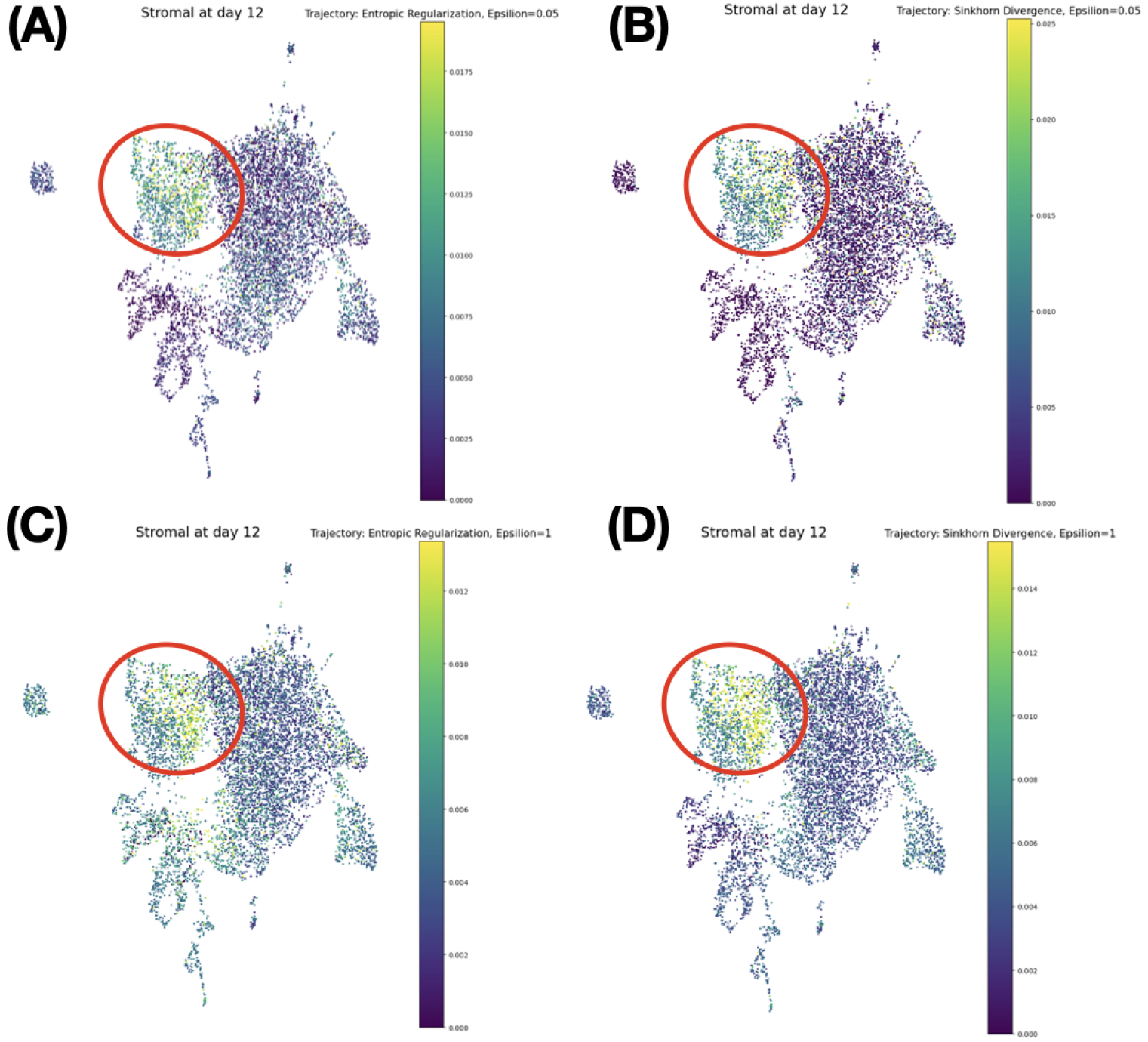
Trajectory Probabilities of stromal cells at Day 12. (A) Entropic transport with *ϵ*=0.05. (B) Sinkhorn transport with *ϵ*=0.05. (C) Entropic transport with *ϵ*=1.0. (D) Sinkhorn transport with *ϵ*=1.0. The clusters of targeted stromal cells are roughly indicated by the red closed loops. Dispersive distributions are evident with entropic regularization whether *ϵ* is 0.05 or 1. When Sinkhorn divergence is applied, the transited distribution is close to the ground truth phenotype with *ϵ* = 0.05, but not with *ϵ* = 1.0.

**Fig. 5.**
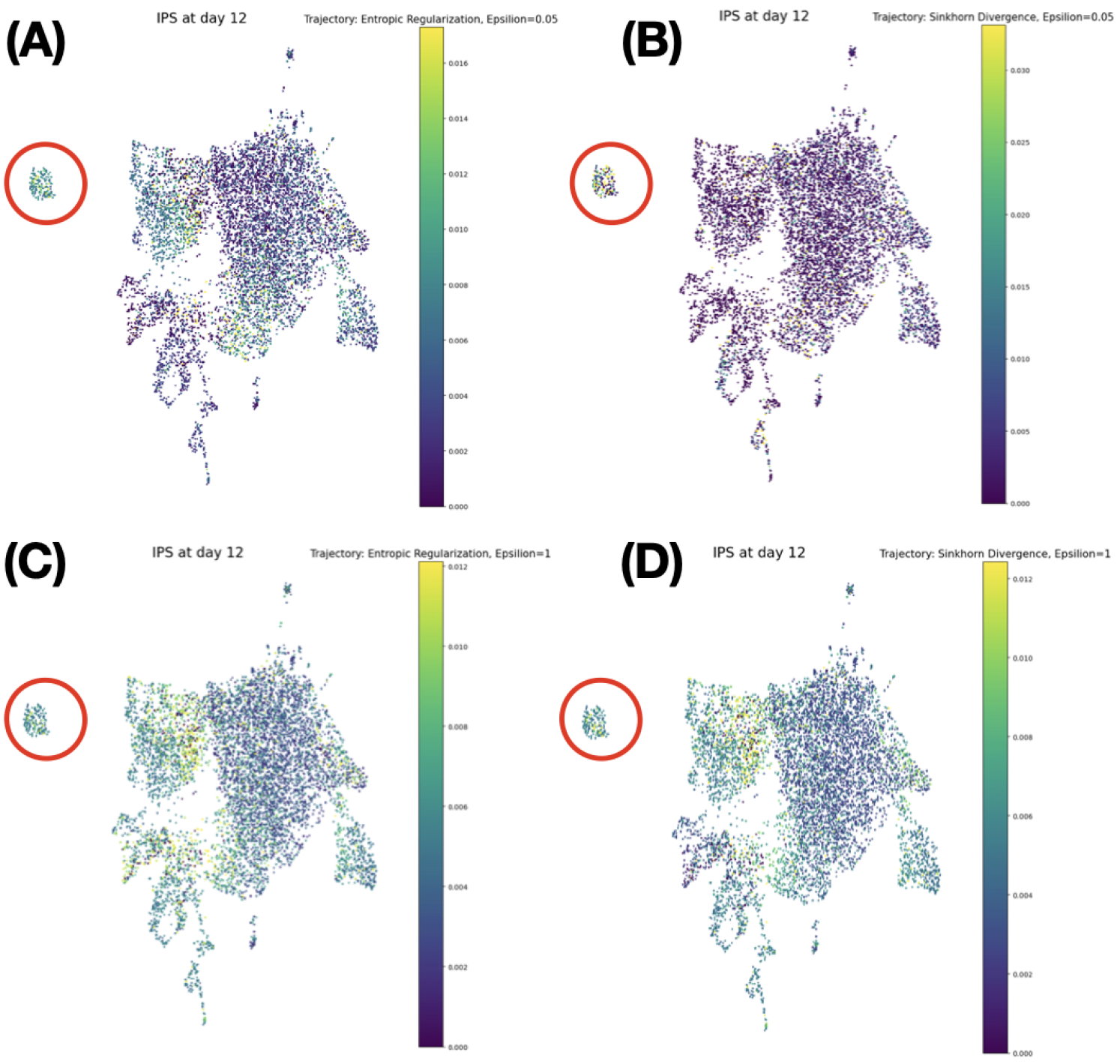
Trajectory Probabilities of iPS cells at Day 12. (A) Entropic transport with *ϵ*=0.05.(B) Sinkhorn transport with *ϵ*=0.05. (C) Entropic transport with *ϵ*=1.0. (D) Sinkhorn transport with *ϵ*=1.0. The clusters of targeted induced pluripotent stem cells are roughly indicated by the red closed loops. Dispersive distributions are evident with entropic regularization whether *ϵ* is 0.05 or 1. When Sinkhorn divergence is applied, the transited distribution is close to the ground truth phenotype with *ϵ* = 0.05, but not with *ϵ* = 1.0.

**Fig. 6.**
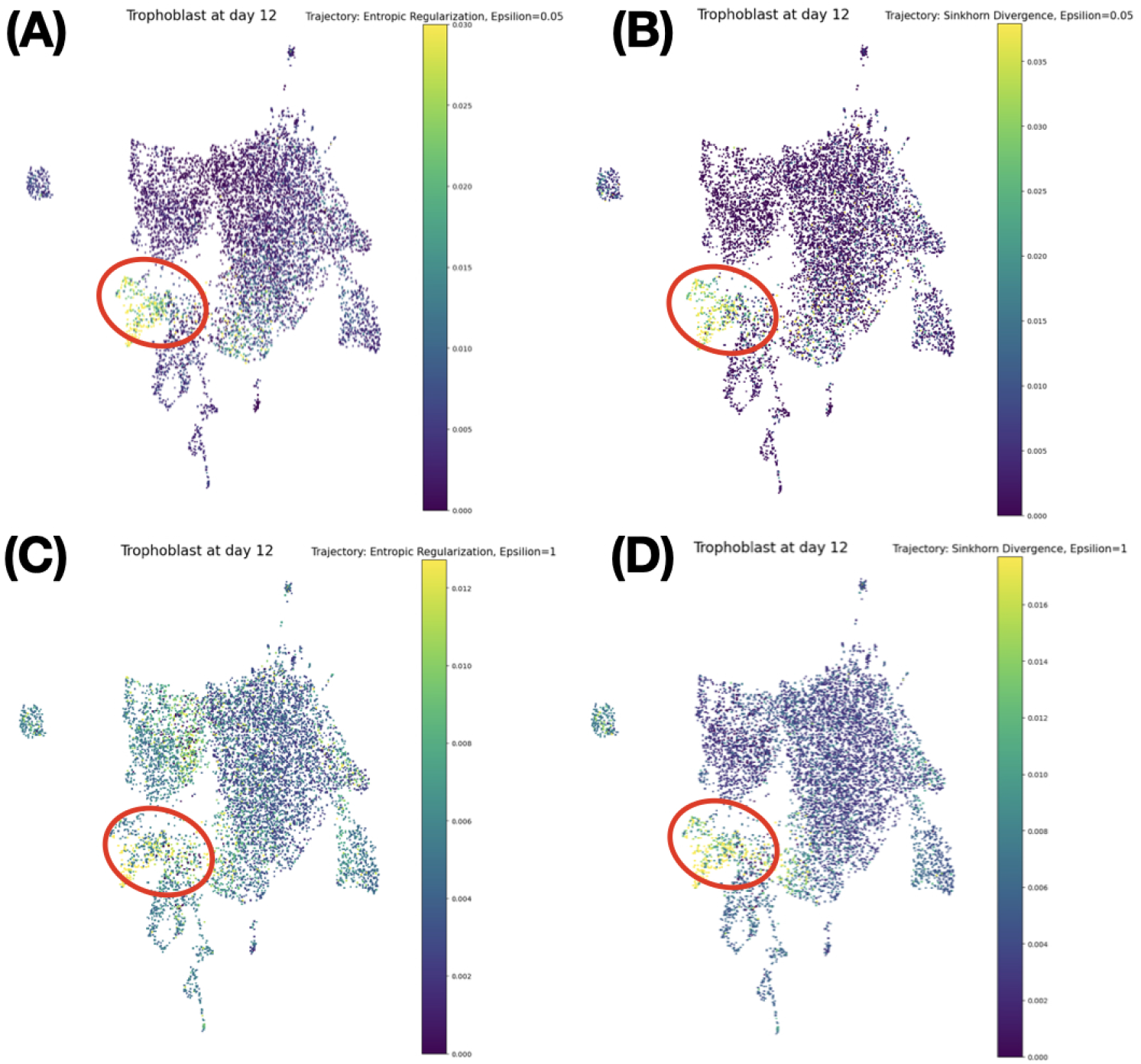
Trajectory Probabilities of trophoblast cells at Day 12. (A) Entropic transport with *ϵ* =0.05. (B) Sinkhorn transport with *ϵ*=0.05. (C) Entropic transport with *ϵ*=1.0. (D) Sinkhorn transport with *ϵ*=1.0. The clusters of targeted trophoblast cells are roughly indicated by the red closed loops. Dispersive distributions are evident with entropic regularization whether *ϵ* is 0.05 or 1. When Sinkhorn divergence is applied, the transited distribution is close to the ground truth phenotype with *ϵ* = 0.05, but not with *ϵ*= 1.0.

**Fig. 7.**
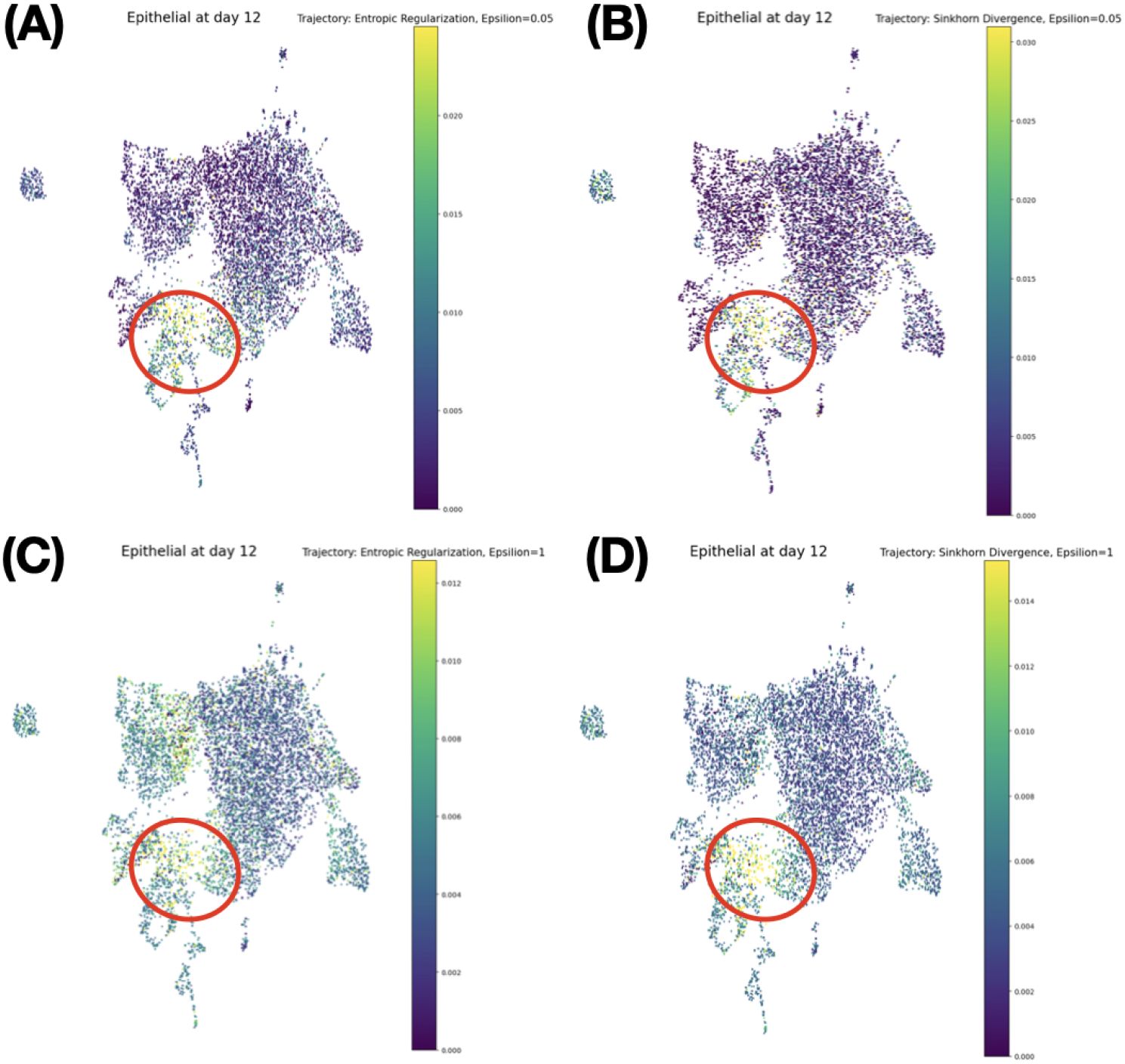
Trajectory Probabilities of Epithelial cells at Day 12. (A) Entropic transport with *ϵ*=0.05. (B) Sinkhorn transport with *ϵ*=0.05. (C) Entropic transport with *ϵ*=1.0. (D) Sinkhorn transport with *ϵ*=1.0. The clusters of targeted epithelial cells are roughly indicated by the red closed loops. Dispersive distributions are evident with entropic regularization whether *ϵ* is 0.05 or 1. When Sinkhorn divergence is applied, the transited distribution is close to the ground truth phenotype with *ϵ* = 0.05, but not with *ϵ* = 1.0.

### Reprogramming to stromal cells

Fig. 4 shows the probability distributions through the trajectory of optimal transport at Day 12 with the targeted phenotype of stromal cells. The red closed loops indicate the subpopulation of ground truth stromal cells qualitatively. Whether Sinkhorn divergence is prescribed or not, *ϵ* = 0.05 conveys a better outcome than *ϵ* = 1.0. When *ϵ* = 0.05, dispersive subpopulations with cell types other than that of stormal cells are observed for the trajectories with entropic regularization without Sinkhorn divergence. As shown in Fig. 3(A) with the convex profile for the Wasserstein distance with the prescription of Sinkhorn divergence, the optimal value of *ϵ* for the best outcome of Sinkhorn divergence is around 0.2. This is within the broad bandwidth for mostly optimal choices of *ϵ* as indicated by the black arrow. In the case of solely applying entropic regularization, the Wasserstein distance monotonically increases in terms of *ϵ*.

### Reprogramming to induced pluripotent stem cells

Fig. 5 shows the probability distributions through the trajectory of optimal transport at Day 12 with the targeted phenotype of induced pluripotent stem cells. The red closed loops specify the islands of subpopulation representing the ground truth iPS cells qualitatively. Whether Sinkhorn divergence is prescribed or not, *ϵ* = 0.05 conveys a better outcome than *ϵ* = 1.0. When *ϵ* = 0.05, dispersive subpopulation is evident for the trajectories with entropic regularization without Sinkhorn divergence. As shown in Fig. 3(B) with the convex profile for the Wasserstein distance with the prescription of Sinkhorn divergence, the optimal value of *ϵ* for the best outcome of Sinkhorn divergence is around 0.3. In the case entropic regularization is solely applied, the Wasserstein distance is the smallest at around *ϵ* = 0.05.

### Reprogramming to trophoblast cells

Fig. 6 shows the probability distributions through the trajectory of optimal transport on Day 12 with the targeted phenotype of trophoblast cells. The red closed loops point to the subpopulation of ground truth trophoblast cells qualitatively. Whether Sinkhorn divergence is prescribed or not, *ϵ* = 0.05 conveys a better outcome than *ϵ* = 1.0. When *ϵ* = 0.05, the measure of dispersed subpopulation seems comparable whether Sinkhorn divergence is applied or not as indicated by Fig. 3(C). The dependency of the Wasserstein distance on *ϵ* shows a convex profile with the prescription of Sinkhorn divergence. The optimal value of for *ϵ* the best outcome of Sinkhorn divergence is around 0.18 with a reasonable tolerance of *ϵ* as indicated by the black arrow. Still, they are worse than the best performance solely with entropic regularization. In the case of solely applying entropic regularization, the Wasserstein distance monotonically increases in terms of *ϵ*. This is the case of Theorem 2 where balancing with Sinkhorn divergence does not show outperformance.

### Reprogramming to epithelial cells

The probability distributions through the trajectory of optimal transport are shown at Day 12 with the targeted phenotype of epithelial cells (Fig. 7). The red closed loops emphasize qualitatively the subpopulation of ground truth epithelial cells. Whether Sinkhorn divergence is prescribed or not, *ϵ* = 0.05 conveys a better outcome than *ϵ* = 1.0. When *ϵ* = 0.05, the measure of dispersed subpopulation seems a little better when Sinkhorn divergence is not applied as indicated by Fig. 3(D). The dependency of the Wasserstein distance on *ϵ* shows a nonlinear profile with the prescription of Sinkhorn divergence. The optimal value of *ϵ* for the best outcome of Sinkhorn divergence is around 0.3. Still, this is worse than the best performance solely with entropic regularization, where the Wasserstein distance monotonically increases in terms of *ϵ*. This is another case of Theorem 2 where balancing with Sinkhorn divergence does not show outperformance.

### Dependency on the time sparsity and population scale

To test the effects of the sparsity of time points from the original reprogramming fibroblast dataset, we have filtered out the time points to 9 (0, 2, 4, 6, 8, 10, 12, 14, 18 days) and 5 (0, 4, 8, 12, 18 days). Also, they are further considered in the effect of cell population size with the main population of *n* = 8000 and the reduced one of *n* = 2000.

In the same way, as done for the original data (Fig. 3), the accuracy of the entropic and Sinkhorn maps in their derived cell lineages with respect to the ground truth cell phenotypes is shown in terms of *ϵ* with the Wasserstein distances (Figs. 8 and 9). For all cell phenotypes, Sinkhorn divergence shows consistent improvement independent of entropic parameter *ϵ*. The black arrows indicate roughly the bandwidh of *ϵ* for mostly optimal choices of entropic regularization in Sinkhorn divergence.

**Fig. 8.**
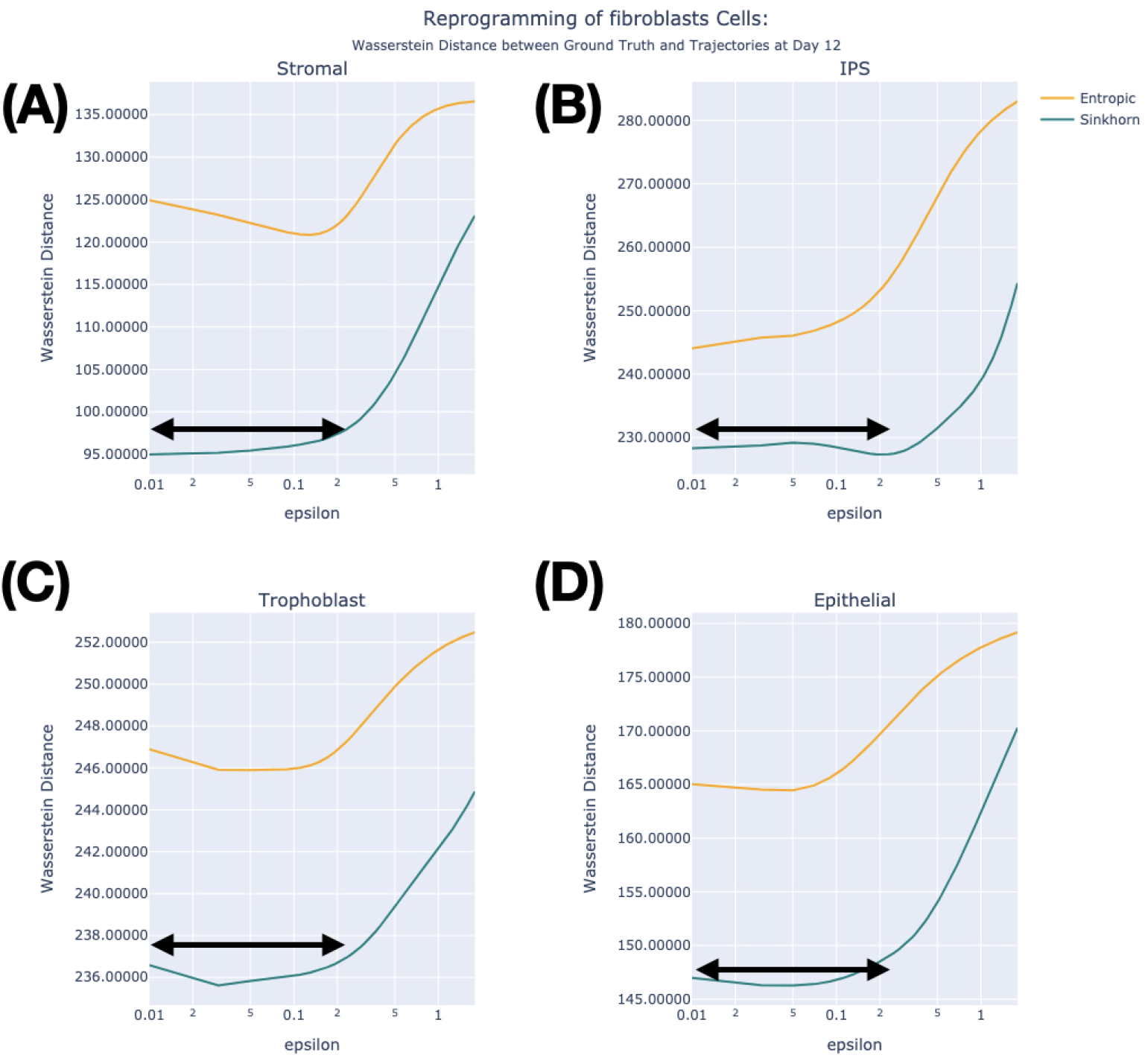
The derived case of fibroblast reprogramming with 9 time points and n = 8000. From the original time points of 36 points for 18 days, 9 time points (0, 2, 4, 6, 8, 10, 12, 14, 18 days) are sampled. The cell population of *n* = 8000 is the same as the original case study (Fig. 3). The accuracy of optimal transport is shown in terms of *ϵ* with Wasserstein distances in probability between ground truth phenotype versus the trajectories from entropic regularization and debiasing Sinkhorn divergence. Each panel is for stromal, iPS, trophoblast, and epithelial phenotypes. For all cell phenotypes, Sinkhorn divergence shows consistent improvement independent of entropic parameter *ϵ*. The black arrows indicate roughly the bandwidth of *ϵ* for mostly optimal choices of entropic regularization for Sinkhorn divergence.

**Fig. 9.**
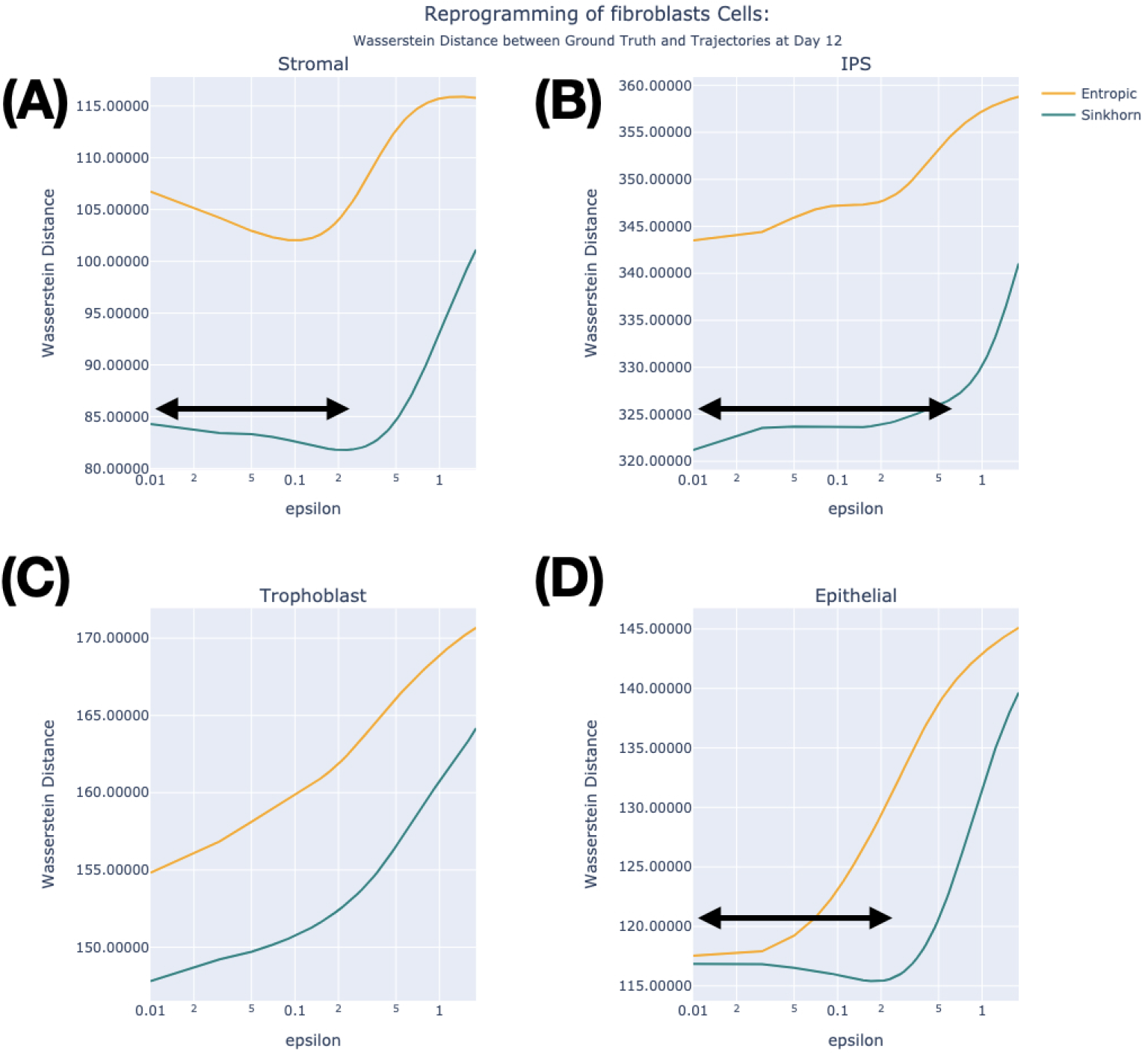
The derived case of fibroblast reprogramming with 9 time points and n = 2000. From the original time points of 36 points for 18 days, 9 time points (0, 2, 4, 6, 8, 10, 12, 14, 18 days) are sampled. The cell population is reduced to *n* = 2000 from the original case study (Fig. 3, *n* = 8000). The accuracy of optimal transport is shown in terms of *ϵ* with Wasserstein distances in probability between ground truth phenotype versus the trajectories from entropic regularization and debiasing Sinkhorn divergence. Each panel is for stromal, iPS, trophoblast, and epithelial phenotypes. For all cell phenotypes, Sinkhorn divergence shows consistent improvement independent of entropic parameter *ϵ*. The black arrows indicate roughly the bandwidth of *epsilon* for mostly optimal choices of entropic regularization for Sinkhorn divergence.

For the cases of 5 time points (four days interval) and *n* = 8000, except for the iPS lineage, Sinkhorn divergence shows a signature of improvement with relevant choices of. When *n* = 2000, except for the trophoblast lineage, Sinkhorn divergence shows a signature of improvement independent of. For trophoblast and epithelium, the best accuracy is from strong entropic regularization. The black arrows roughly indicate the tolerated bandwidth of for mostly optimal choices of entropic regularization in Sinkhorn divergence (Figs. 10 and 11).

**Fig. 10.**
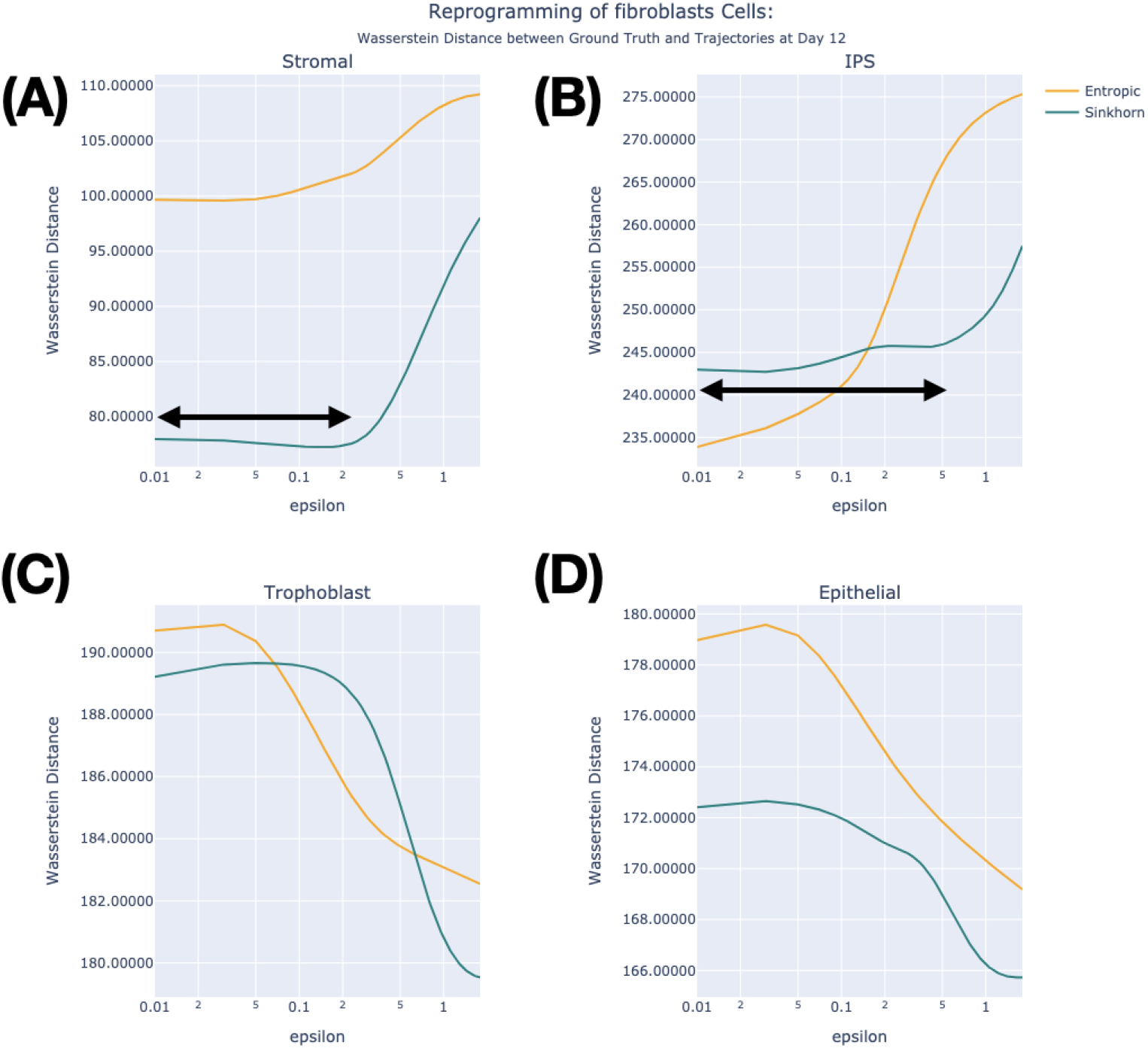
The derived case of fibroblast reprogramming with 5 time points and n = 8000. From the original time points of 36 points for 18 days, 5 time points (0, 4, 8, 12, 18 days) are sampled. The cell population size *n* = 8000 is the same as the original case study (Fig. 3). The accuracy of optimal transport is shown in terms of *ϵ* with the Wasserstein distances in probability between ground truth phenotype versus the trajectories from entropic regularization and debiasing Sinkhorn divergence. Each panel is for stromal, iPS, trophoblast, and epithelial phenotypes. Except for iPS lineage, Sinkhorn divergence shows a signature of improvement with relevant choices of *ϵ*. The black arrows roughly indicate the tolerated bandwidth of *ϵ* for mostly optimal choices of entropic regularization for Sinkhorn divergence.

**Fig. 11.**
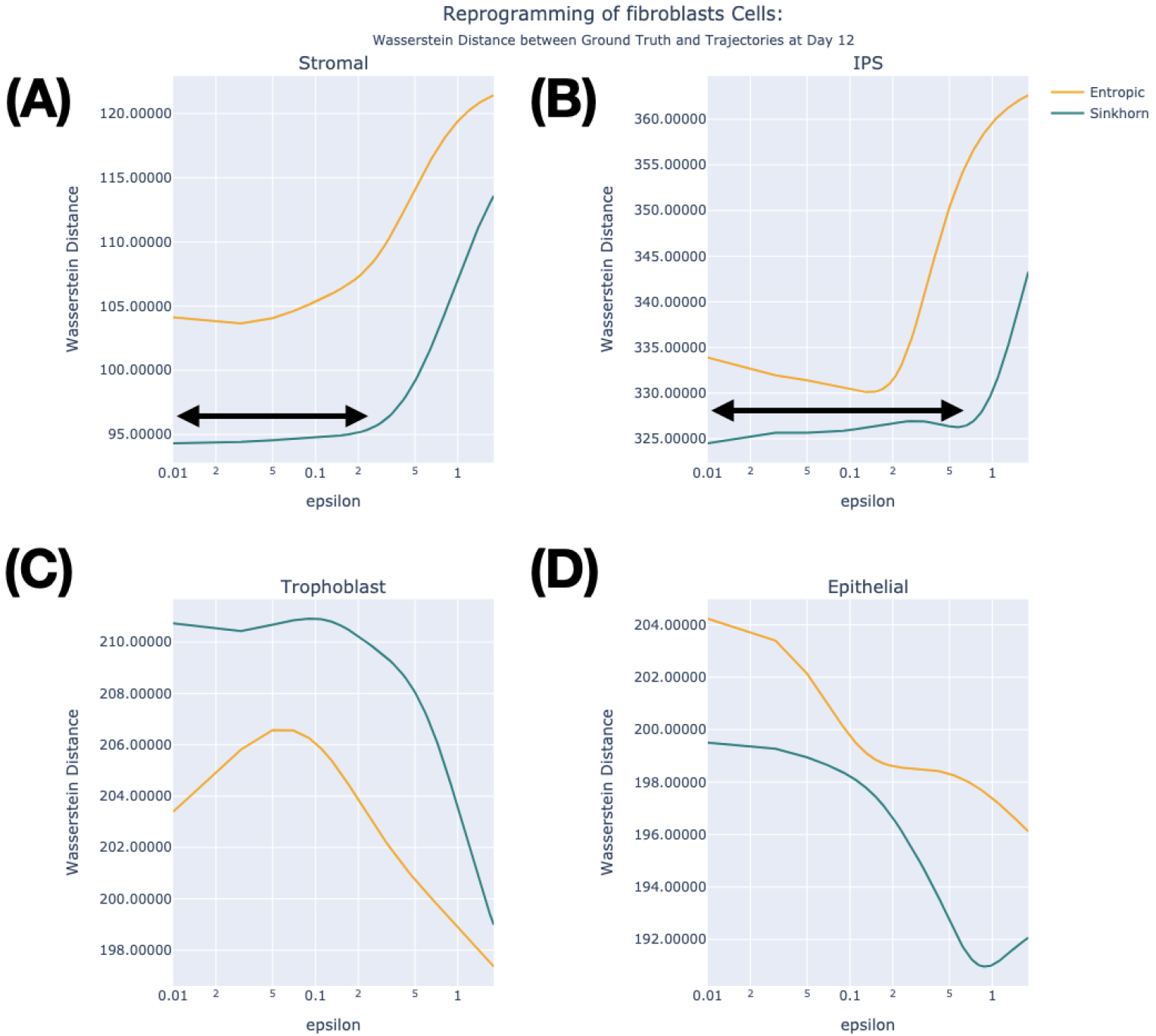
The derived case of fibroblast reprogramming with 5 time points and n = 8000. From the original time points of 36 points for 18 days, 9 time points (0, 4, 8, 12, 18 days) are sampled. The cell population size *n* = 2000 is a reduced one from the original case study (Fig. 3). The accuracy of optimal transport is shown in terms of *ϵ* with Wasserstein distances in probability between ground truth phenotype versus the trajectories from entropic regularization and debiasing Sinkhorn divergence. Each panel is for stromal, iPS, trophoblast, and epithelial phenotypes. Except for trophoblast lineage, Sinkhorn divergence shows a signature of improvement independent of *ϵ*. For trophoblast and epithelium, the best accuracy is from strong entropic regularization. The black arrows roughly indicate the tolerated bandwidth of *ϵ* for mostly optimal choices of entropic regularization for Sinkhorn divergence.

### 4.2 Case Study 2: Stratification of epidermal cells in morphogenesis

Two embryonic time points in morphogenesis (E12.5 and E17.5) are used for single-cell RNA sequencing experiments on wild-type epidermal cells [2]. From a dataset of 1,350 cells collected over an 18-day period, the modified Waddington-OT algorithm with Sinkhorn divergence is applied to a subset of around 950 cells. The cells at E12.5 are transited to E17.5 by optimal transport in the expectation of programming them to differentiating suprabasal, hair follicular, interfollicular epidermis cells as shown in Fig. 2(B). In the same way as Case Study 1, one track is through the trajectories with entropic regularization. The other track is through the trajectories with entropic regularization balanced by Sinkhorn divergence. For each cell type, the influence of entropic regularization is determined with the variation of. Accordingly, we have quantified the Wasserstein distance between the probability distributions of the ground truth from experimental data and the derived phenotypes from the trajectories of optimal transport in terms of (Fig. 12). The probability distributions through the trajectories of hair follicular and differentiating suprabasal cells at E17.5 are shown in Figures 13 and 14, respectively. Those four subpanels show the combination of *ϵ* of 0.05 or 1.0 and the entropic regularization with and without Sinkhorn divergence. Consistent to Theorem 2, for differentiating suprabasal cell types, small entropic parameters do not guarantee the outperformance of Sinkhorn divergence.

**Fig. 12.**
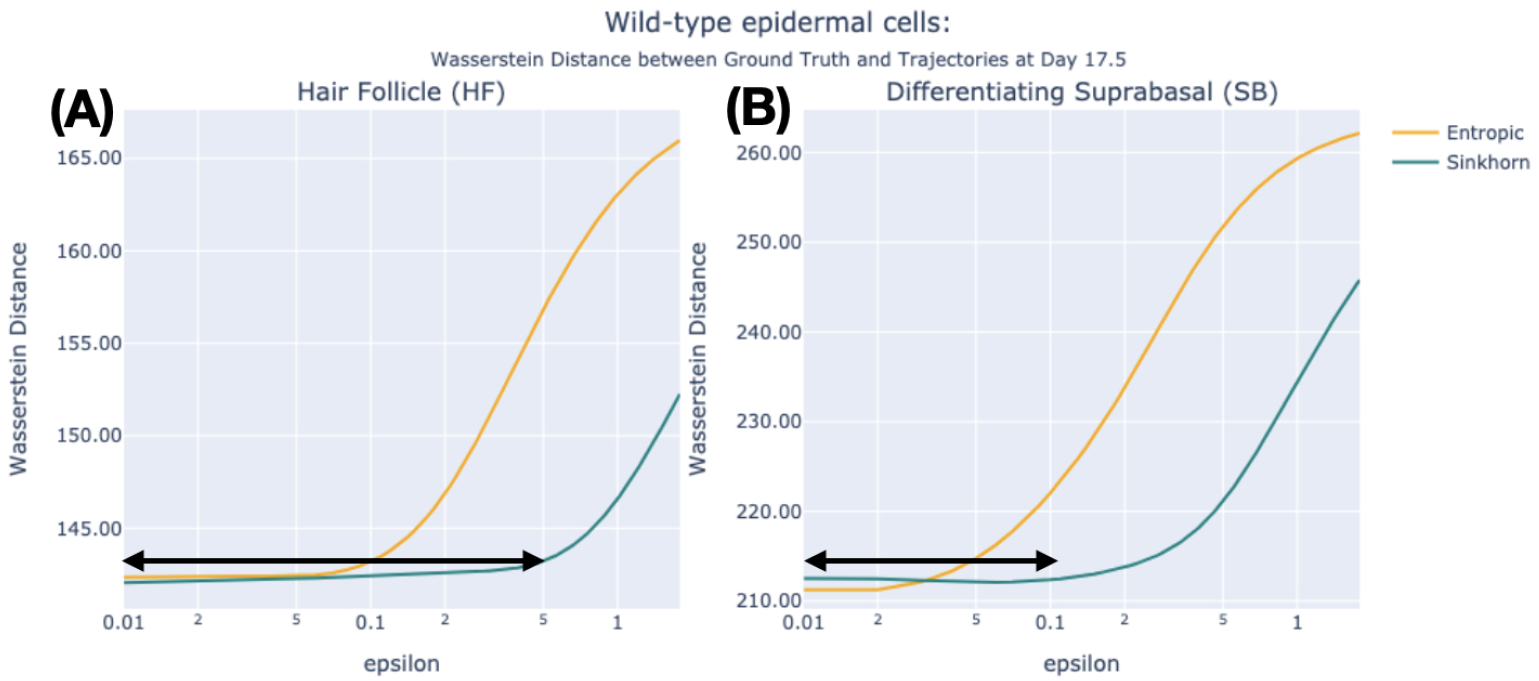
The accuracy of optimal transport for epidermal morphogenesis. Wasserstein distances between the probability distributions of the ground truth phenotype and the trajectories from the entropic map and Sinkhorn map are plotted in terms of the entropic parameter *ϵ* in the range from 0.01 to 2. The best performance is when *ϵ* is small and Sinkhorn divergence does not show significant benefits of debiasing. The black arrows roughly indicate the tolerated bandwidth of *ϵ* for mostly optimal choices of entropic regularization in Sinkhorn divergence.

**Fig. 13.**
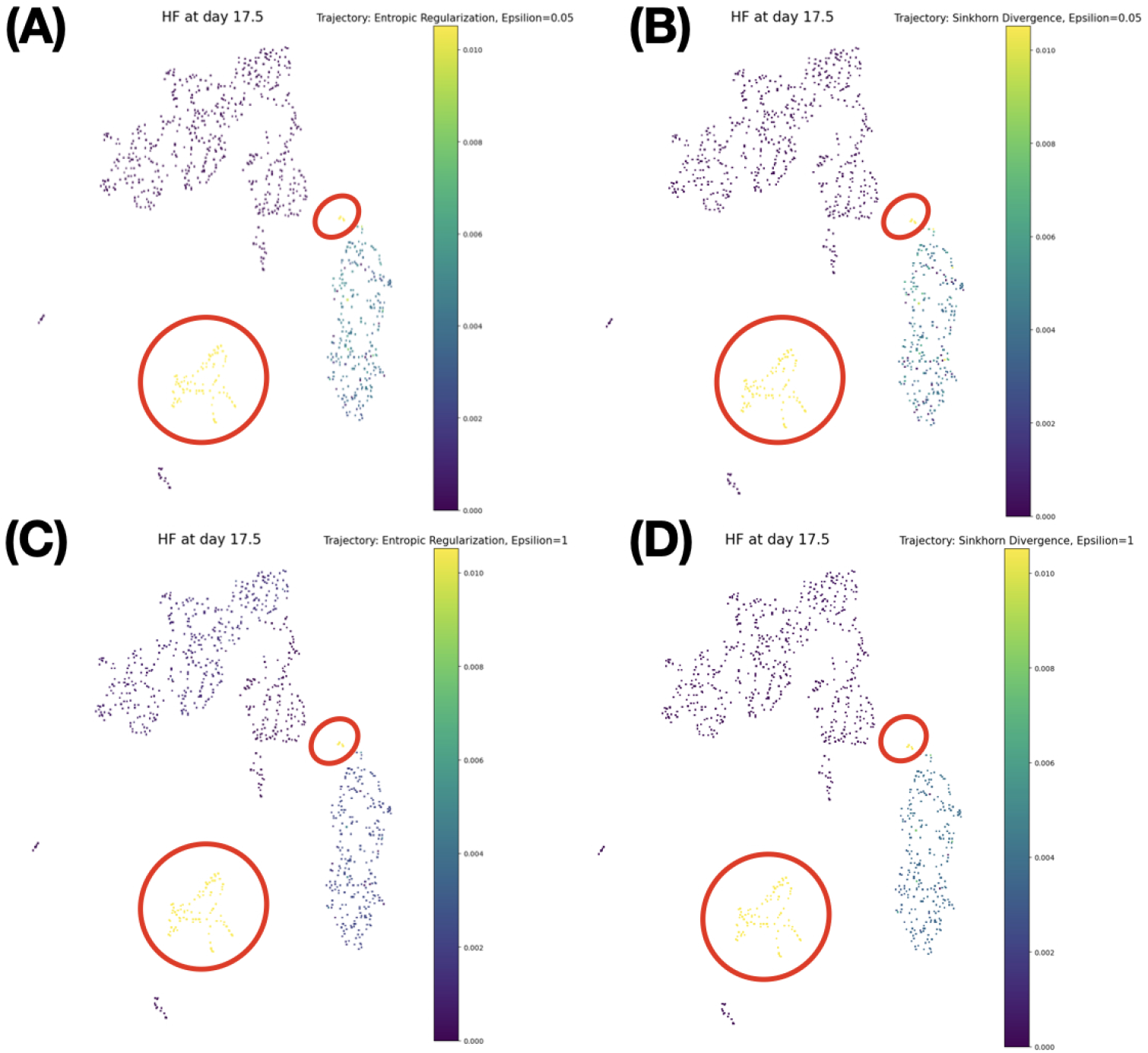
Hair Follicle (HF) Trajectories Probabilities at E17.5. (A) Entropic transport with *ϵ*=0.05. (B) Sinkhorn transport with *ϵ*=0.05. (C) Entropic transport with *ϵ*=1.0. (D) Sinkhorn transport with *ϵ*=1.0. The clusters of targeted hair follicle cells are roughly indicated by the red closed loops.

**Fig. 14.**
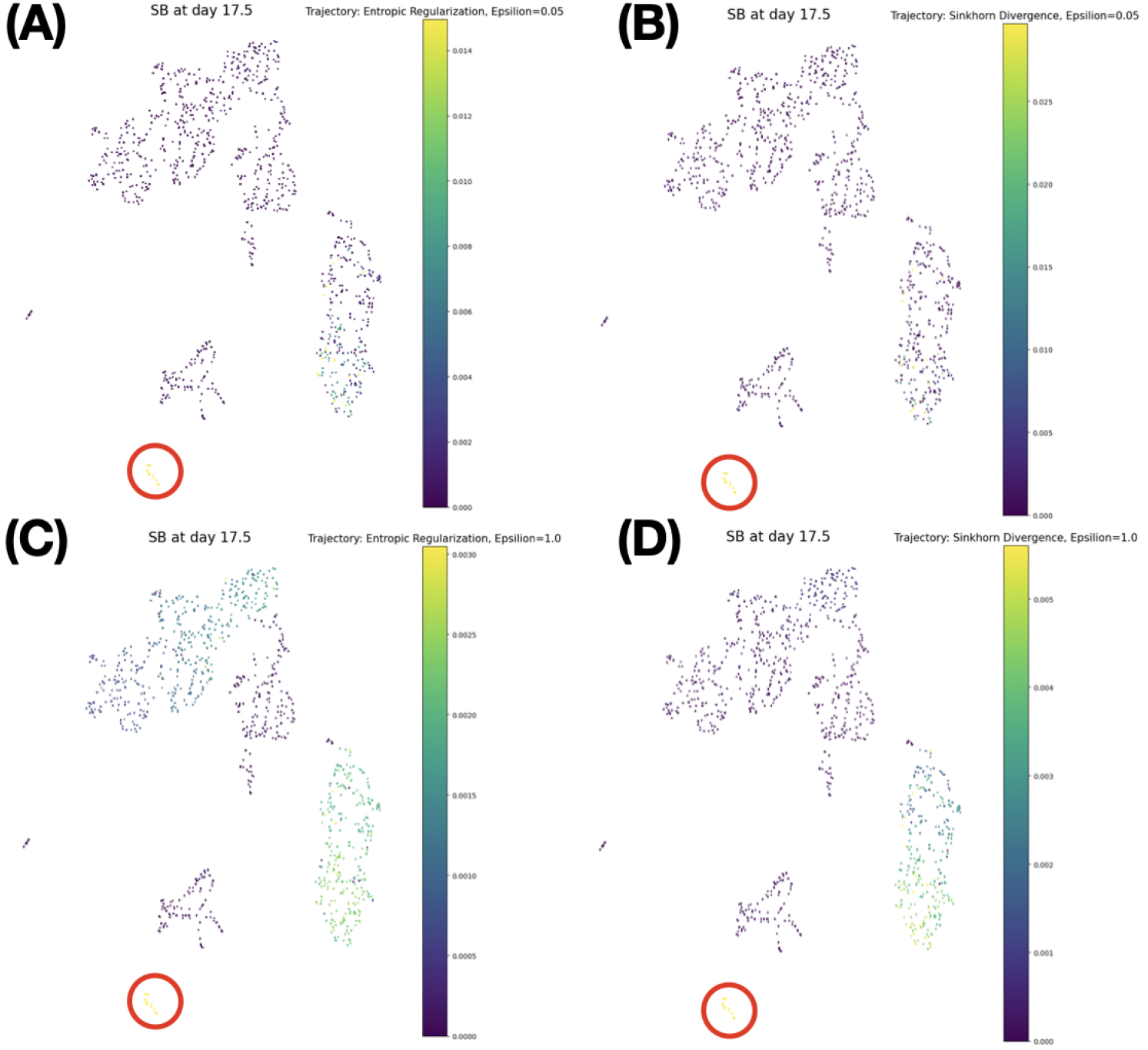
Suprabasal (SB)Trajectories Probabilities at Embryonic Day 17.5. (A) Entropic transport with *ϵ* =0.05. (B) Sinkhorn transport with *ϵ* =0.05. (C) Entropic transport with *ϵ* =1.0. (D) Sinkhorn transport with *ϵ* =1.0. The clusters of targeted epithelial cells are roughly indicated by the red closed loops. Dispersive distributions are evident with entropic regularization whether *ϵ* is 0.05 or 1. When Sinkhorn divergence is applied, the transited distribution is close to the ground truth phenotype with *ϵ* = 0.05, but not with *ϵ* = 1.0.

### Programming to hair follicle cells

Fig. 13 shows the probability distributions through the trajectory of optimal transport at E17.5 targeted to hair follicle cells. The red closed loops point to the subpopulation of ground truth hair follicle cells qualitatively. Whether Sinkhorn divergence is prescribed or not, small values of *ϵ* do provide a bandwidth of plateau with small Wasserstein distances and the most optimal accuracy as shown in Fig. 12(A). There is no significant benefit of prescribing Sinkhorn divergence.

### Programming to suprabasal cells

The probability distributions through the trajectory of optimal transport are shown at E17.5 targeted to suprabasal cells in Fig. 14. The red closed loops specify qualitatively the subpopulation of ground truth suprabasal cells. When *ϵ* is 0.05, the measure of dispersed subpopulation seems a little better when Sinkhorn divergence is applied as indicated by Fig. 14. When *ϵ* is 1.0, dispersive subpopulations are observed for the trajectories whether Sinkhorn divergence is prescribed or not. Similar to the case of transiting hair follicle cells, independent of Sinkhorn divergence, small values of *ϵ* do provide a bandwidth of plateau with small Wasserstein distances and the most optimal accuracy as shown in Fig. 12(B). There is no significant benefit to prescribing Sinkhorn divergence.

## 5 Conclusion

The existing framework of optimal transport for single cells with entropic regularization, so-called Waddington-OT, is balanced with Sinkhorn divergence. Even though the underlying mathematical structure of Sinkhorn divergence is symmetric and balanced, the issue of generating coherent trajectories toward targeted cell fates and phenotypes needs empirical experiments. With two showcases of datasets where one is dense and the other is sparse in time points, indeed, we have shown the non-trivial aspects of prescribing Sinkhorn divergence and determining the optimal values for entropic regularization, where time points and sample size are critical factors. Also, we observed that Sinkhorn divergence makes the choice of entropic parameter tolerated in broad bandwidths in many cases. It remains how to overcome the sparsity in the time course data. One possibility is to integrate the Eulerian formalism of Kantorvich and Lagrangian formalism of dynamic optimal transport as an iterative scheme filling in the sparse time points from inferred transport models within the constrained optimization. This is a class of Eulerian-Lagrangian interaction in optimal transport. In addition, cell-to-cell interaction is missing part of the current formulation of modeling cell dynamics and possibly one source of the present gap between the ground truth and the outcome of the trajectories from optimal transport. In the Lagrangian formalism, cell-to-cell interaction can be well represented with interacting potentials.

Furthermore, when spatial single-cell analysis is considered, Gromov-Wasserstein metrics can be employed and the associated debiasing algorithm will be open avenues to explore. This article lays a foundation for these future works.

## 6 Acknowledgement

J. Cooper, C. Young, and P. Lee were partially supported by NIH Data Science and NIMHD grant, 3U54MD013376-04S3.

